# Asynchronous mixing of kidney progenitor cells potentiates nephrogenesis in organoids

**DOI:** 10.1101/761916

**Authors:** Ashwani K. Gupta, Prasenjit Sarkar, Xinchao Pan, Thomas Carroll, Leif Oxburgh

**Author notes:** Author for correspondence:. Mailing address: The Rogosin Institute, 310 East 67^th^ Street, New York, NY 10065. Feinberg School of Medicine, Northwestern University, Chicago, Illinois, USA.

## Abstract

Recent years have seen rapid advances in directed differentiation of human pluripotent stem cells (PSCs) to kidney cells. However, a fundamental difficulty in emulating kidney tissue formation is that kidney development is iterative. Recent studies argue that the human nephron forms through gradual contribution of nephron progenitor cells whose differentiation fates depend on the time at which they are recruited. We show that the majority of PSC-derived nephron progenitor cells differentiated in a short wave in organoid formation and to improve fidelity of PSC-derived organoids, we emulated the asynchronous mix found in the fetal kidney by combining cells differentiated at different times in the same organoid. Asynchronous mixing promoted nephrogenesis, and lineage marking data showed that proximal and distal nephron components preferentially derive from cell populations differentiated at distinct times. When engrafted under the kidney capsule these heterochronic organoids were vascularized and displayed essential features of kidney tissue. Micro-CT and injection of a circulating vascular marker demonstrated that engrafted kidney tissue was connected to the systemic circulation by 2 weeks after engraftment. Proximal tubule glucose uptake was confirmed using intravenous injection of fluorescent dextran. Despite these promising measures of graft function, overgrowth of stromal cells prevented long-term study, and we propose that this is a technical feature of the engraftment procedure rather than a specific shortcoming of the directed differentiation because kidney organoids derived from primary cells and whole embryonic kidneys develop the same stromal overgrowth when engrafted under the kidney capsule.

## INTRODUCTION

Kidney injury is a common consequence of several prevalent conditions such as type 2 diabetes, cardiovascular disease and systemic lupus erythematosus. In addition, drug toxicity, ischemia, and genetic mutations cause primary kidney injury. It is estimated that approximately 15% of adults have reduced kidney function (https://www.usrds.org), and these individuals are predisposed to end stage kidney disease, in which function is impaired to the point that basic physiological processes cannot be maintained. For these approximately 750,000 patients in the USA (https://www.usrds.org), dialysis is generally the first step, but the small molecule filtration and volume adjustment accomplished in routine dialysis does not compensate for all of the functions of the kidney tissue that have been lost. This is one important explanation for the discrepancy in survival between dialysis and kidney transplantation (Kaballo et al., 2018), although contributing factors are multifactorial. Currently, over 100,000 patients are waiting for a kidney transplant in the USA (https://optn.transplant.hrsa.gov) and based on statistics from 2018, it is anticipated that approximately 20% will receive an organ this year. Each day 13 people die waiting for a kidney transplant and there is a pressing need to seek new strategies to increase the availability of tissue.

Although we have known since the 1950s that mammalian kidneys can be cultured *in vitro* (Grobstein, 1953), attempts to grow transplantable tissue from primary cells and embryonic organs have met with limited success. Recent years have seen rapid advances in regenerative medicine following the discovery that adult somatic cells can be programmed back to induced pluripotent stem cells (iPSCs) from which their differentiation can be directed along any organ lineage. Theoretically, this provides two major advantages. First, it may be possible to recreate the complex interaction of cell types required for organ formation because multiple cell types can be differentiated simultaneously through directed differentiation. Second, tissues derived from iPSCs are autologous with the individual from which the iPSCs were reprogrammed, which should ensure minimal transplant rejection.

Procedures for the directed differentiation of human pluripotent stem cells (PSCs) to kidney cells have been developed (Morizane et al., 2015; Taguchi et al., 2014; Takasato et al., 2015), and in these procedures, pluripotent stem cells are first differentiated to kidney progenitor cells, then triggered to epithelialize using a pulse of the GSK3 inhibitor CHIR, which is thought to mimic the natural epithelialization stimulus Wnt9b. This single epithelialization pulse differs from the process of differentiation in the developing kidney, where cells at numerous stages of differentiation coexist within the organ. Recent work has argued for a model in which the nephron forms through association of differentiating epithelial cells with cells that derive from the progenitor population, suggesting a requirement for asynchronous cell populations for tissue formation. We show that emulating this process in vitro through mixing of cells at distinct stages of differentiation promotes nephrogenesis, and we provide lineage marking data proving that proximal and distal nephron components preferentially derive from different cell populations. When engrafted under the kidney capsule these heterochronic organoids are vascularized and display essential features of kidney tissue. However, overgrowth of stromal cells is an obstacle that prevents long-term study of these grafts, and we propose that this problem is a technical feature of the engraftment procedure rather than a specific shortcoming of the directed differentiation procedure as kidney organoids derived from primary cells and whole embryonic organs develop the same stromal overgrowth when engrafted under the kidney capsule.

## RESULTS

### Strategy for formation of kidney organoid tissue for engraftment

Directed differentiation of PSCs generates a mix of kidney progenitor cells that is predicted to represent the repertoire of cell types within the mesoderm that gives rise to the nephron, interstitium and vasculature of the fetal kidney. We therefore reasoned that culturing these cells in the organotypic conditions developed for culture of the rodent kidney (Saxen, 1987) would be a logical starting point for the formation of new human kidney tissue from PSCs. After performing the 9 day directed differentiation procedure for WTC11 iPSCs or H9 human embryonic stem cells (ESCs) as described by (Morizane et al., 2015), we dispersed the cells and aggregated them by resuspending the cells in a low volume to form a dense slurry and pipetting carefully into a droplet on the hydrophobic surface of a polycarbonate filter. At this point culture conditions were switched to those developed for primary nephron progenitor cells (NPCs) (Brown et al., 2015); APEL 2 medium supplemented with protein free hybridoma medium, BMP7, FGF9, and heparin. In preparatory experiments, we found that the addition of BMP7, FGF9 and heparin promote the viability of cultured cells generated by directed differentiation (data not shown) and we determined that these factors could be removed from the medium after the first 4 days of aggregate culture to reduce cost (Fig. 1A). The mix of cells resulting from directed differentiation is predicted to contain either no or only a very small number of progenitors for the collecting duct system (Islam and Nishinakamura, 2019), which is known to serve as the inducer of nephron epithelium differentiation (Carroll et al., 2005). In prior experiments we have provided the Wnt stimulus that induces mesenchyme-to-epithelium transition through treatment with the small molecule GSK3 inhibitor CHIR 99021 (Brown et al., 2015; Gupta et al., 2019). However, in a series of titrations aimed at determining the dose of CHIR to use to initiate directed differentiation of WTC11 iPSCs to kidney progenitors, spontaneous epithelial differentiation can be seen at day 9 of the directed differentiation, prior to detachment of cells and organoid formation (Fig. S1), and we therefore reasoned that cells may not require CHIR 99021 treatment for epithelial differentiation in aggregate culture. CHIR 99021 is a potent inhibitor of GSK3 and is expected to have multiple effects other than activating Wnt/β-catenin signaling. Thus, omitting it from our differentiation protocol for tissue engraftment would be highly beneficial as it would reduce the scope for off-target effects. We found that kidney progenitor cells derived from directed differentiation of PSCs did indeed form epithelia following aggregation without CHIR treatment in our conditions (Fig. 1B, C). However, the epithelialization process does not appear efficient as large areas of undifferentiated cells can be seen in aggregates. For engraftment, we reasoned that it is important to convert the vast majority of cells in the aggregate to epithelium in order to avoid out-competition by undifferentiated cell types over the several-week period required for vascularization/perfusion, and we therefore developed the aggregate culture method further to enhance mesenchyme-to-epithelium transition.

**Figure 1:**
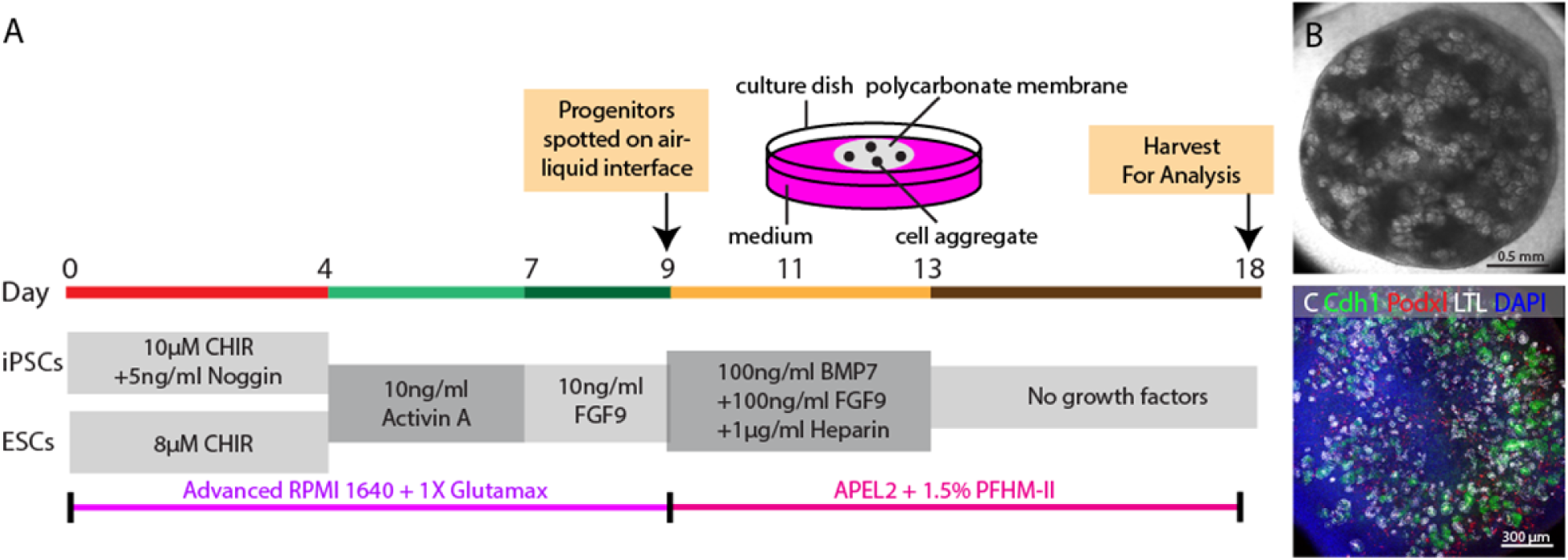
Generation of kidney organoids through directed differentiation. **(A)** Schematic diagram showing directed differentiation of hPSCs into kidney progenitors through Morizane protocol and further aggregation to a kidney organoid. **(B)** Representative stereo microscope image of kidney organoid on day 18. **(C)** Immunofluorescence staining for Podocytes (Podxl), Proximal tubule (LTL) and distal tubule (Cdh1).

Nephron formation in the kidney occurs in waves and undifferentiated NPCs are located immediately adjacent to epithelializing nephrons. Recent studies of human fetal kidney development suggest that there is gradual contribution of NPCs to the forming nephron, and that the differentiation fate of cells is dependent on when they are recruited into the nascent structure; time-dependent fate acquisition (Lindstrom et al., 2018a). To establish a culture system in which NPCs could be added to epithelializing nephrons over time, we mixed newly differentiated cells with cells that had been aggregated on polycarbonate filters to allow formation of epithelia. To ascertain at which stages of differentiation it might be possible to improve epithelialization by this heterochronic recombination, we staggered directed differentiation protocols so that we could mix newly differentiated cells with cells that had been aggregated in organotypic conditions for 1, 2, or 3 days (Fig. S2). We found that recombination of newly differentiated kidney progenitor cells with cells that had been aggregated and cultured for 2 days generated organoids with the most tightly packed epithelial structures (Fig. 2A-C). To quantify this effect, we compared the degree of staining for molecular markers of podocyte (podocalyxin), proximal tubule glycoproteins (lotus lectin), and distal tubule (cadherin 1) between organoids derived from an aggregate of a single population of cells from directed differentiation versus organoids from a heterochronic mix (Fig. 2D). Compared with aggregates cultured from only the first batch of directed differentiation cells or from only the second batch of directed differentiation cells, aggregates generated by heterochronic mixing displayed approximately double the number of structures stained for each molecular marker. On close examination, aggregates derived by heterochronic mixing contained proximal tubule cells (Fig. 2E), distal tubule cells (Fig. 2F), connecting segment or collecting duct (Fig. 2G), podocytes (Fig. 2H), endothelial cells (Fig. 2H), and stromal cells (Fig. 2I). Interestingly, the network of CD31-expressing presumptive endothelial cells in organoids from heterochronic mixing is more extensive and complex than that seen in organoids generated from a single directed differentiation population (Fig. S3).

**Figure 2:**
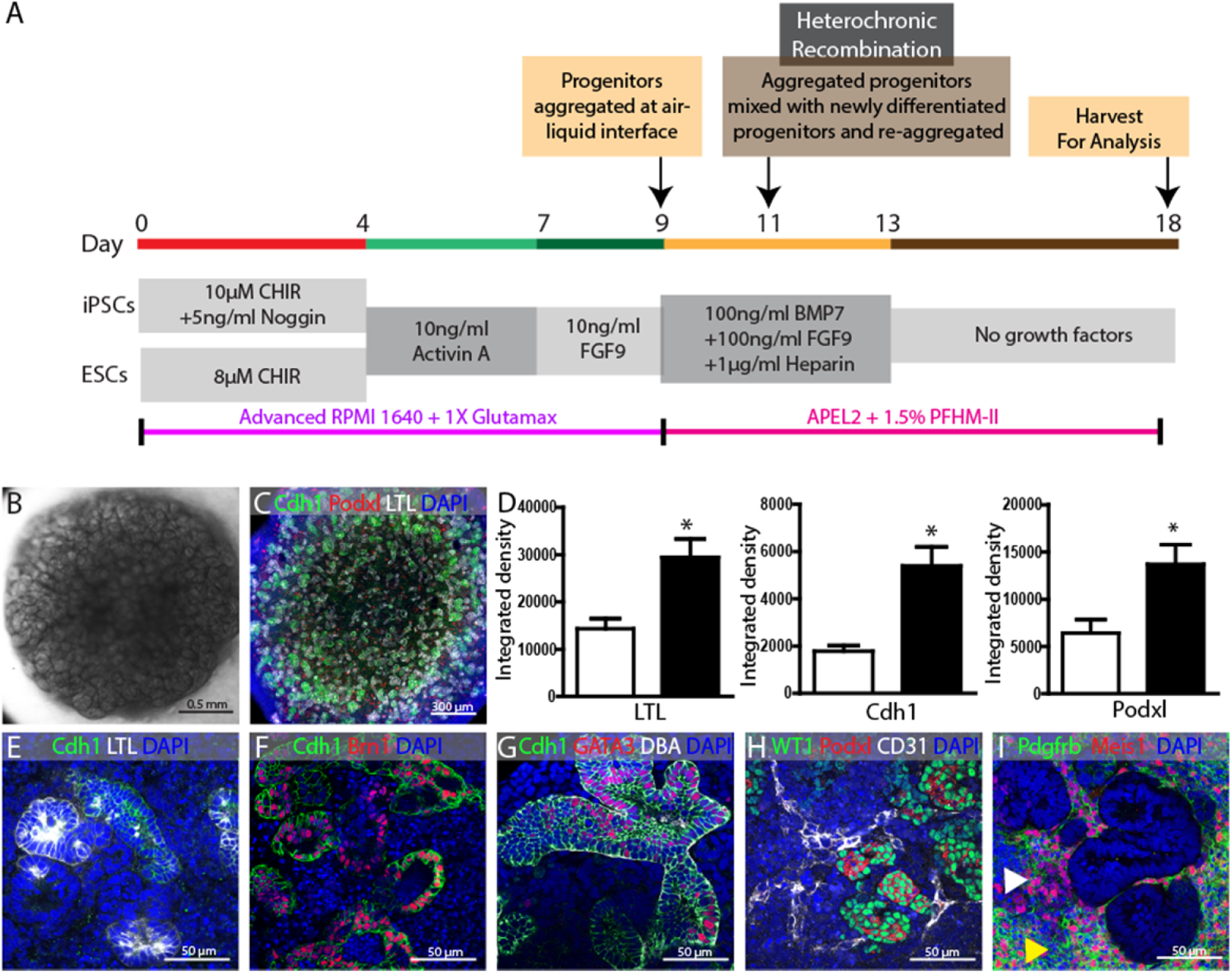
Generation of kidney organoids through heterochronic recombination. **(A)** Schematic illustration of protocol for heterochronic mixing, in which 2 directed differentiations are staggered by 2 days so that the first batch may be aggregated and cultured for 2 days before the second, newly differentiated, batch is added. **(B)** Representative stereo microscope image of kidney organoid generated through heterochronic recombination. **(C)** Immunofluorescence staining for Podocytes (Podxl), Proximal tubule (LTL) and distal tubule (Cdh1) in the organoid. **(D)** Integrated density for LTL, Cdh1, and Podxl in organoids generated from single batch (open bar) and heterochronic mixing (solid bar). Values calculated from three independent experiments. Statistical significance was determined by unpaired t-test and expressed as mean±SEM; *p<0.05. **(E-I)** Representative high magnification immunofluorescence images of organoids derived from heterochronic recombination showing LTL^+^ Cdh1^-^ proximal tubule, Brn1^+^ Cdh1^+^ distal tubule, CDH1^+^ GATA3^+^ DBA^+^ connecting tubule or collecting duct, Podxl^+^ WT1^+^ Podocytes, CD31^+^ endothelial network, Pdgfrb^+^ Meis1^-^ Pericytes (yellow arrow head) and Pdgfrb^+^ Meis1^+^ stromal cells (white arrow head).

In summary, our protocol efficiently generates organoids with tightly packed nephron epithelia and endothelial networks that are suitable for engraftment without the need for stimulation of organoid epithelialization by treatment with CHIR 99021 or other Wnt stimulators that may cause off-target effects.

### Contributions of the two heterochronic cell batches to nephron and interstitial cell types

To determine if the two batches of cells used for heterochronic recombination contribute equivalently to the different cell types in the epithelialized aggregate, we performed a series of experiments in which we combined H9 human embryonic stem cells with H9 cells that have been modified to express a fluorescently tagged H2B histone subunit under the control of the ubiquitous CAG enhancer-promoter. Fluorescently tagged H9 cells (H9-FP) were either incorporated as the first batch with unlabeled cells as the second batch (Fig. 3A) or as the second batch, with unlabeled batch 1 cells (Fig. 3B). H9-FP cells introduced in either batch 1 or batch 2 contribute to podocyte (Fig. 3C,D), proximal tubule epithelium (Fig. 3E,F), distal tubule epithelium (Fig. 3G,H), and interstitial cells (Fig. 3I,J). However, quantification reveals that batch 2 cells preferentially contribute to the podocyte (Fig. 3K) and proximal tubule (Fig. 3L), while batch 1 cells preferentially contribute to distal epithelium (Fig. 3M) and interstitial cells (Fig. 3N). This observation is consistent with microanatomical characterization of fetal kidneys showing a time-dependent fate acquisition of NPCs in which the podocyte population is added late in the formation of the nascent nephron (Lindstrom et al., 2018a). To understand if the basis of the synergistic effect seen upon addition of newly differentiated cells could be replenishment of NPCs that are depleted early in the process of epithelialization, we quantified cells labeled with the NPC markers SIX2 and WT1 and the marker of early NPC differentiation LHX1. Flow cytometry analysis was performed on both heterochronic batches: newly differentiated cells and cells that had been aggregated for 2 days. Consistent with the original report (Morizane et al., 2015), the newly differentiated cell mix contains approximately 90% SIX2+ cells (Fig. 3O). This proportion is reduced to 70% after 2 days of aggregate culture. Conversely, the proportion of LHX1+ cells (Fig. 3P) increases from 35% in newly differentiated cells to 86% after 2 days of aggregate culture indicating that the majority of cells have undergone differentiation. In the fetal kidney, some SIX2 expression is maintained in differentiating cells of the early nephron (Lindstrom et al., 2018b), which may explain why SIX2 expression is maintained at 70% while 86% of cells express the differentiation marker LHX1. Interestingly, the independent NPC marker WT1 (Fig. 3Q) is expressed in 57% of newly differentiated cells but marks only 26% of cells after 2 days of aggregate culture. In summary, 2 days of aggregate culture result in widespread differentiation of LHX1+ cells and depletion of SIX2+/LHX1-NPCs, supporting the notion that improved differentiation of the heterochronic mix compared to a single batch is due to replenishment of the NPC population.

**Figure 3:**
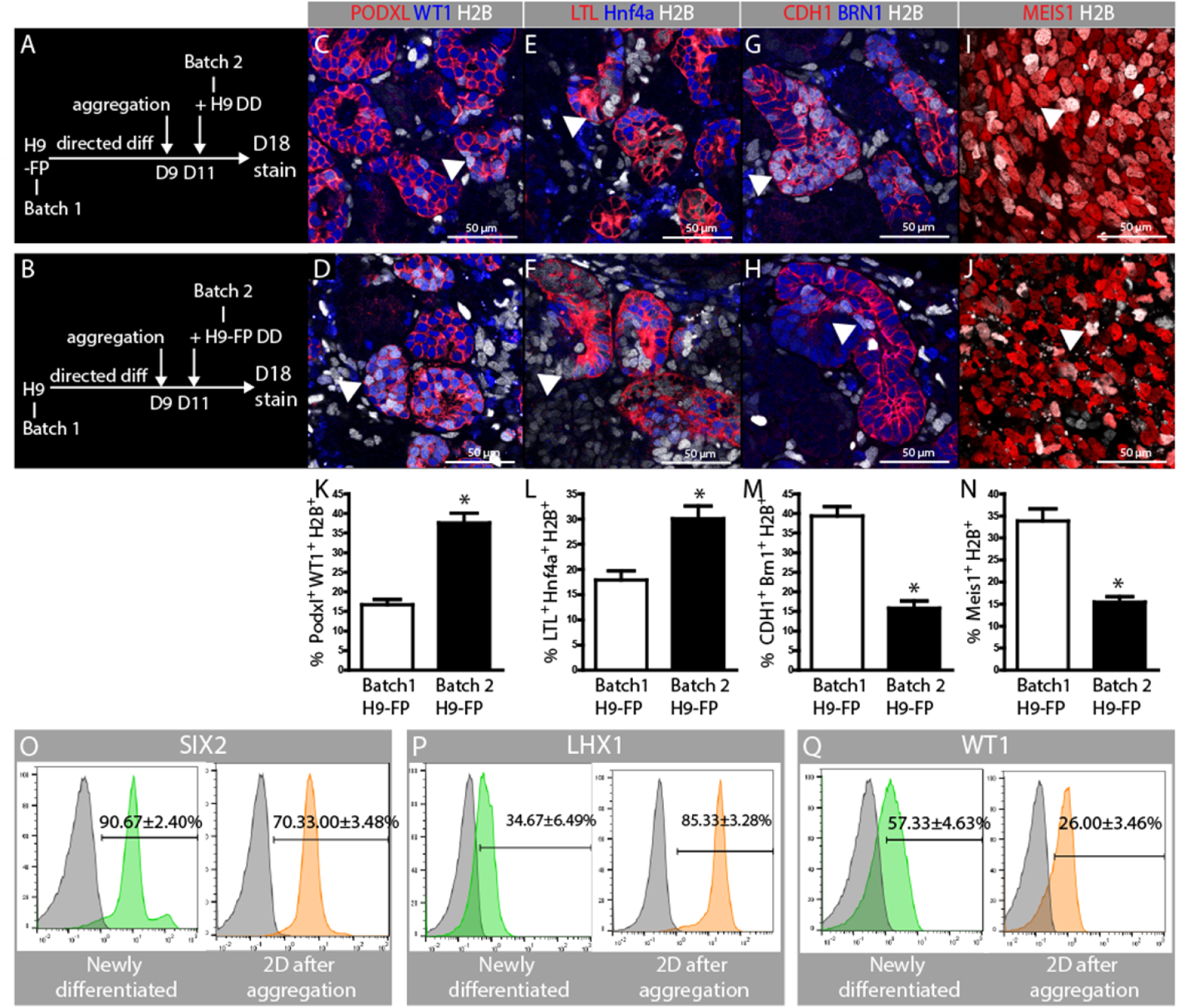
Contribution of heterochronic cell batches to nephron segments and interstitial cells. **(A-B)** Experimental plans showing two strategies for heterochronic recombination of H9 and H9-FP cells, which constitutively express. **(A)** H9-FP cells were aggregated following directed differentiation and newly differentiated H9 cells were added 2 days thereafter. In **(B)**, H9 cells were aggregated following directed differentiation and H9-FP cells were added 2 days thereafter. **(C-D)** H2B-marked cell contribution to PODXL+/WT1+ podocytes. **(E-F)** H2B-marked contribution to LTL+/HNF4a+ proximal tubules. **(G-H)** H2B-marked contribution to CDH1+/BRN1+ distal tubules. **(I-J)** H2B-marked contribution to MEIS1+ interstitial cells. Arrowheads in each of the panels indicate H2B-marked cells that have contributed to the labeled cell populations. **(K-N)** shows quantification of contribution of H2B marked cells introduced in the batch 1 versus those introduced in batch 2 to each of the labeled cell populations. Cells in three random fields with differentiated structures were counted in each organoid. Values calculated from three independent experiments, statistical significance was determined by unpaired t-test and expressed as mean±SEM. *p<0.05. (O-Q) Flow cytometric comparison of the proportion of cells staining for SIX2, LHX1 and WT1 in newly differentiated cultures (green graphs) versus cells that have been aggregated and cultured as organoids for 2 days (orange graphs). Values expressed as mean±SEM from three independent experiments.

### Engrafted kidney organoids are vascularized and form both nephron and interstitial cell types

To determine if kidney tissue derived from heterochronic recombination forms perfused tissue in vivo, we engrafted organoids under the kidney capsules of severely immunocompromised NSG mice. The scheme for organoid differentiation and engraftment is shown in Figure 4A. Twenty organoids of 1-1.5mm diameter and 0.25-0.5mm thickness were engrafted into a single subcapsular site in each animal, and animals were sacrificed 3 weeks after engraftment. Extensive growth of engrafted tissue was seen (Fig. 4B-D), and kidneys were vibratome sectioned for whole mount molecular marker analysis. Staining with the endothelial marker CD31 revealed widespread vascularization of the graft, emanating from the host kidney (Fig. 4E). Higher magnification imaging reveals patent vessels connecting with clusters of WT1/Podxl-labeled presumptive podocytes (Fig. 4F). Glomeruli with characteristic morphology including complex capillary structures and Bowman’s space are found throughout the graft (Fig. 4G,H). Staining for the extracellular matrix proteins collagen I, collagen IV and laminin reveals basement membrane morphology similar to glomeruli in the host (Figure S4); HNF4a and LTL label proximal tubule (Fig. 4I,J,K); cadherin1, Tamm Horsfall protein, and BRN1 mark distal tubule segments (Fig. 4J,K,L); GATA3, DBA, and KRT8 mark extreme distal tubule segments or collecting duct (Fig. 4M,N); Renin is expressed in subsets of cells at the base of glomeruli indicating differentiation of the juxtaglomerular apparatus (Fig. 4O). In summary, marker analysis showed that the cardinal cell types of the kidney are present in the graft, and that the tissue is vascularized. To ascertain which of the cells in grafts derive from human and which derive from mouse, we stained tissue for human nuclear antigen (Fig. 4P-U). Counterstain for the cell types listed above revealed that all cell types that differentiate in the subcapsular space derive from the human graft, with the exception of the CD31 and endomucin-expressing endothelium (Fig. 4T,U), which appears to be exclusively derived from the host.

**Figure 4:**
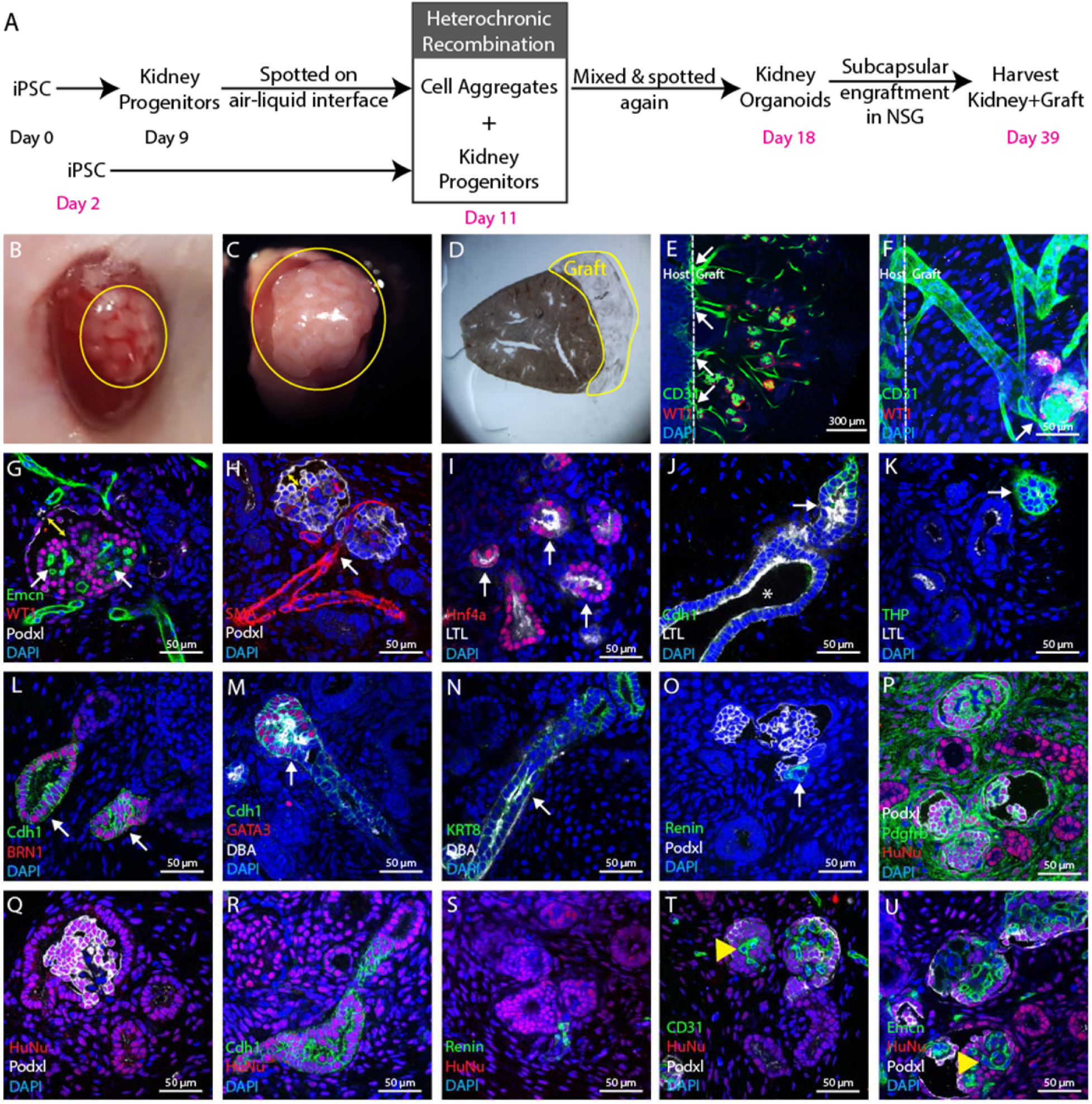
In vivo maturation of kidney organoids. **(A)** Schematic diagrams showing generation of kidney organoids and their subcapsular engraftment in mice kidney. **(B)** Image showing kidney organoids immediately after engraftment. **(C)** Three weeks after engraftment organoids grows extensively in a big tissue. **(D)** Stereo microscope image showing vibratome section of mouse kidney with graft for further analysis. **(E)** Low magnification immunofluorescence image showing CD31+ endothelial network coming from host kidney widespread in the graft. **(F)** High magnification immunofluorescence image showing these CD31+ endothelial networks are connecting with Podxl^+^ WT1^+^ presumptive glomeruli. **(G-H)** Immunofluorescence images showing characteristic morphology of glomerular structure including Podxl^+^ WT1^+^ podocytes, Emcn^+^ capillary tuft and bowman’s space (yellow double arrow head). These glomerular structures also shows direct connection with SMA^+^ vascular network. **(I)** Immunofluorescence image showing Hnf4a^+^ and LTL^+^ proximal tubule and **(J)** this LTL^+^ Cdh1^-^ proximal tubule segmented in LTL^-^ Cdh1^+^ distal tubule with lumen (white arrow head). **(K)** Immunofluorescence image showing THP^+^ loop of Henle and **(L)** Brn1^+^ Cdh1^+^ distal tubule. **(M-N)** Immunofluorescence image showing Cdh1^+^ GATA3^+^ DBA^+^ connecting tubule or collecting duct connected with Cdh1^+^ GATA3^-^ DBA^-^ distal segment of nephron. In addition, immunofluorescence images showing KRT^+^ DBA^+^ extreme distal segment or collecting duct. **(O)** Further, immunofluorescence image showing Renin^+^ juxtaglomerular cells at the base of Podxl^+^ glomerular structure. We didn’t observe Renin^+^ cells in organoid before engraftment (data not shown) which is another confirmation of organoid maturation in vivo. **(P-S)** Immunofluorescence image showing Pdgfrb^+^ mesangial cells surrounded by Podxl^+^ Podocytes inside presumptive glomerular structures. Further immunofluorescence images showing differentiated Pdgfrb^+^ mesangial or interstitial cells, Podxl^+^ podocytes, Cdh1^+^ epithelial structures and Renin^+^ cells were derived from human cells (HuNu^+^) but **(T-U)** Emcn^+^/CD31^+^ endothelial cells were derived from mouse (HuNu^-^, yellow arrow head). Images shown here are representative and were derived from n=6 NSG mice.

### Micro computed tomography reveals perfusion 2 weeks after engraftment

Having ascertained that organoids derived from heterochronic recombination of human kidney progenitors from directed differentiation are vascularized and maintained in vivo, we wanted to develop a method with which we could track the perfusion of grafts without having to sacrifice mice. This would enable longitudinal studies of graft retention and open up possibilities to monitor performance of the graft following drug treatment. Attempts to use MRI showed some promise but the long scan times required for imaging together with the high cost deterred us from pursuing this strategy. Instead, we used traditional X-ray technology with an iodinated contrast agent. To enable three dimensional imaging, we developed settings for micro-computed tomography. Animals were imaged 3 times at 1-week intervals following engraftment using the procedure outlined in figure 5A. As expected, only calcified structures and kidneys are radio-opaque in animals following administration of contrast agent; note the absence of signal in the adrenal at the pole of the kidney (Fig. 5B). Although contrast agent circulates through soft tissues surrounding the kidneys, it does not accumulate there and the concentration is insufficient to generate a signal at the settings used. Accumulation of sufficient quantities of iodinated compound in the graft to generate signal would signify that the contrast agent is being cleared through the tissue and would provide both a confirmation that the tissue is perfused, and some measure of function. Interestingly, subcapsular grafts are not radio-opaque after 1 week (Fig. 5B), indicating that they are either not yet adequately perfused, or that they are not sufficiently functional to accumulate iodinated compound. By 2 weeks after surgery (Fig. 5C), the graft has become diffusely radio-opaque confirming vascularization. By 3 weeks (Fig. 5D) the morphology of the contrast image has changed and there are regions of intense accumulation of iodinated compound within the graft, indicating functional compartmentalization within the graft.

**Figure 5:**
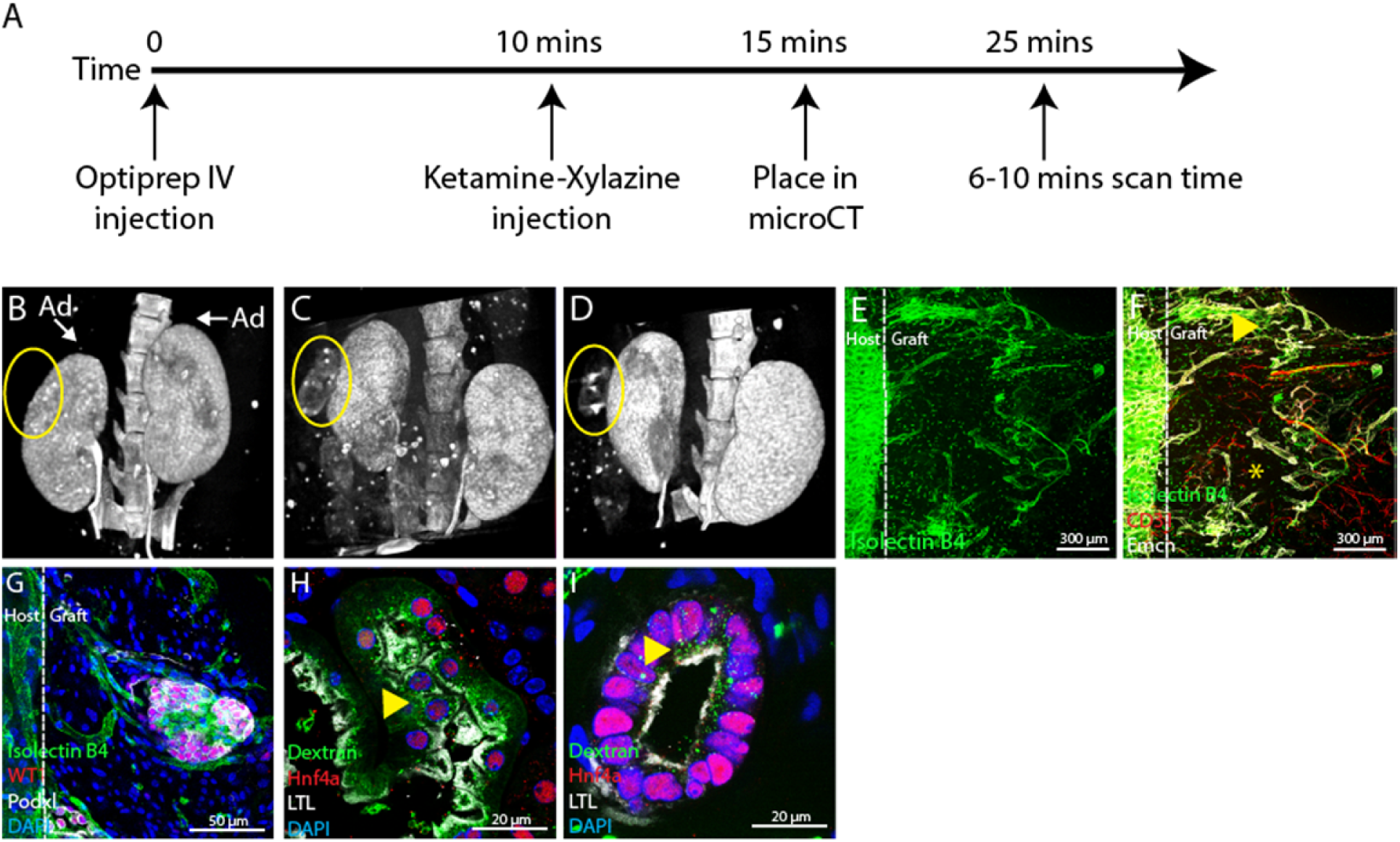
Micro computed (μ-CT) tomography on live mice and functional characterization of the graft. **(A)** Schematic strategy for micro computed tomography; Optiprep contrast agent was intravenously injected in mice that were anesthetized and scanned 25 minutes after injection. Following the 6-10 minute scan, mice were allowed to recover from anesthesia and re-scanned at 2 and 3 weeks after engraftment. **(B)** μ-CT image 7 days after PSC organoid engraftment showing contrast agent accumulation only in the kidney of the host and not in the graft (circled). Note that there is no contrast agent accumulation in any abdominal soft tissues adjacent to the kidney, for example the adrenals (Ad). **(C)** μ-CT 14 days after engraftment shows diffuse accumulation of contrast agent in the subcapsular graft (circled) as well as in the host kidneys. **(D)** μ-CT 21 days after engraftment shows punctate contrast agent accumulation in the graft (circled) as well as accumulation in the host kidney. **(E)** Fluorescence image showing circulation of FITC-labeled isolectin B4 in a graft 3 weeks after subcapsular engraftment. FITC-labeled isolectin B4 was intravenously injected in the orbital sinus, and circulation into the graft implies that it is connected to the systemic circulation. **(F)** Co-staining of FITC-isolectin B4 perfused graft with the endothelial markers endomucin (Emcn) and CD31. Immunofluorescence image showing graft is compartmentalized into highly vascularized regions (yellow arrow head) and poorly vascularized regions (yellow asterisk), consistent with the μ-CT image 3 weeks after engraftment. **(G)** High power image of FITC-isolectin B4 costained with the podocyte markers WT1 and podocalyxin (Podxl) shows that presumptive glomeruli in the graft are connected to the systemic circulation 3 weeks after engraftment. **(H-I)** Fluorescently labeled dextran was injected into the orbital sinus 30 minutes before sacrifice to determine if proximal tubule epithelial cells of the graft take it up in vesicles. **(H)** Fluorescent dextran vesicle uptake (yellow arrowhead) in epithelial cells of the host that stain with the proximal tubule markers HNF4a and LTL. **(I)** Fluorescent dextran vesicle uptake (yellow arrowhead) in epithelial cells of the graft that stain with the proximal tubule markers HNF4a and LTL. Mice for this experiment were harvested 3 weeks after engraftment. All images shown in this figure are representative of 3 engrafted mice per experiment.

To confirm connection of the graft vasculature to the systemic circulation, we administered a retro-orbital injection of FITC-labeled Griffonia Simplificata Isolectin B4 (FITC-IB4) to host animals immediately prior to sacrifice. FITC-IB4 binds glycoproteins on the luminal surface of endothelial cells, and can thus be used to confirm that systemic blood is circulating through graft vasculature. Whole mount imaging of vibratome sections of engrafted tissue revealed extensive networks of labeled vessels (Fig. 5E). Interestingly, co-staining with the endothelial markers CD31 and endomucin reveal that the graft is compartmentalized into the highly vascularized and poorly vascularized regions (Fig. 5F). Within vascularized regions, FITC-IB4 stained vasculature is intimately associated with WT1/PODXL-labeled clusters of presumptive podocytes (Fig. 5G), supporting systemic circulation through glomeruli in the graft. Systemic circulation through glomeruli and evidence of distended Bowman’s capsules (Fig. 4G,H,P,Q) indicates that blood is being filtered in the graft tissue. To test whether filtration might be occurring, we administered retroorbital injections of fluorescently labeled dextran to hosts and monitored uptake in the proximal tubules of the graft. Dextran (glucose) in the systemic circulation passes over the glomerular filtration barrier and is actively reclaimed by epithelial cells of the proximal tubule. An example of a proximal tubule from the host kidney is shown in figure 5G; note the accumulation of FITC-dextran in vesicles. Similarly, HNF4a and LTL labeled proximal tubule epithelial cells within the graft show uptake FITC-dextran (Fig. 5H), although the size of vesicles appears more modest than that seen in the host kidney. Based on these findings, we conclude that there is circulation of systemic blood through the vasculature of the graft, that this blood is filtered, and that the proximal tubule is sufficiently differentiated to actively reclaim glucose. Thus, the engrafted human kidney tissue derived from directed differentiation shows hallmark signs of function, providing an important proof of principle and motivating further development of this technology with the aim of functional testing in disease models. One issue that complicates the use of these grafts for functional testing is the overabundance of stromal cells, which are highly proliferative and in time may out-compete functional graft tissue.

### Proliferation of stroma in grafted organoids from primary cells suggests that environmental factors rather than incomplete iPSC/ESC differentiation are the primary cause of stromal overabundance

One important question in understanding the overabundance of stromal cells in grafts of PSC-derived kidney tissue is if the subcapsular environment in the adult kidney promotes stromal proliferation in kidney tissue or if this is a problem specific to tissue generated from PSCs. To answer this question, we engrafted E15.5 embryonic kidneys subcapsularly in adult NOD SCID kidneys (Fig. 6A). Vigorous growth of the embryonic implant could be seen 1 month after engraftment (Fig. 6B), with abundant vascularized glomeruli (Fig. 6C,D). Immunostaining with the stromal marker PDGFRβ revealed widespread areas of stromal cells between glomeruli, suggesting that factors associated with subcapsular engraftment promote stromal abundance. To confirm using a primary cell system more closely analogous to PSC-derived organoids, we used the nephrogenic zone cell isolation technique (Blank et al., 2009) to generate organoids composed of primary E17.5 nephrogenic zone cells using a similar aggregation and organotypic culture method as was used for PSC-derived directed differentiation cultures. Organoids derived from NZCs are morphologically very similar to those derived from PSCs (Fig. 6F). NZC organoids were subcapsularly engrafted in adult NSG mice and harvested after 3 weeks. At this point, extensive glomerulogenesis was noted (Fig. 6G), and the tissue showed abundant investment of CD31+ endothelial cells (Fig. 6H). However, staining for PDGFRβ again showed an over-abundance of stromal cells (Fig. 6I), providing more evidence that the adult subcapsular environment promotes these cells in grafts of primary kidney tissue.

**Figure 6:**
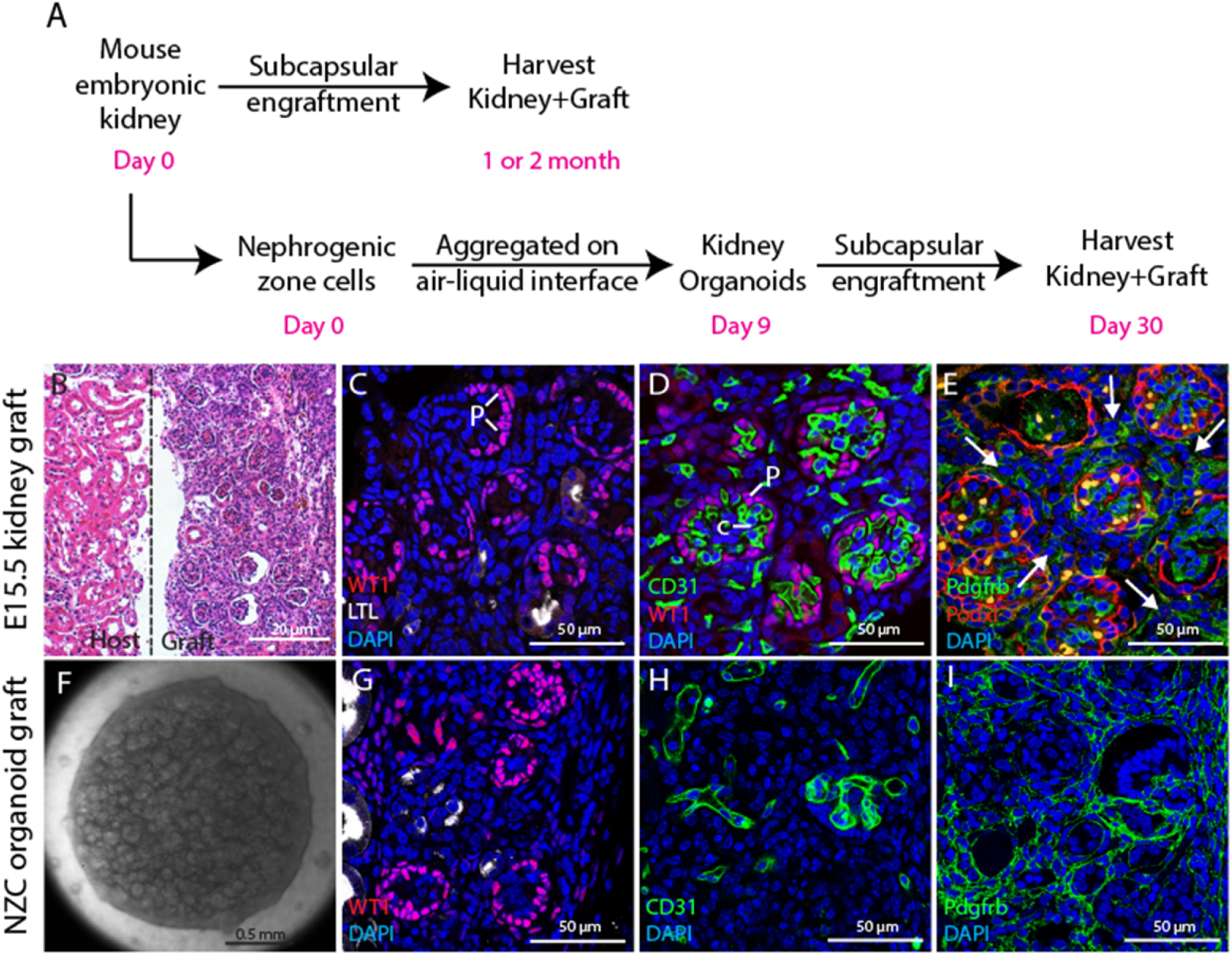
Subcapsular engraftment of embryonic kidney tissue results in stromal over-abundance. **(A)** NSG mice were engrafted with embryonic kidney tissue using two distinct strategies: E15.5 whole embryonic kidneys were subcapsularly engrafted and harvested after one or two months, or nephrogenic zone cells were isolated from E17.5 kidneys, aggregated to organoids, subcapsularly engrafted and harvested after 2 weeks. **(B-E)** Tissue from E15.5 kidney graft. **(B)** H&E staining reveals extensive graft growth. **(C)** WT1 staining of podocytes shows abundant glomeruli and LTL staining for proximal tubules that these are sparse in the graft tissue. **(D)** CD31 staining of endothelial cells shows that glomeruli are vascularized. **(E)** Podocalyxin (Podxl) staining outlines glomeruli and PDGFRβ staining reveals abundant stromal cells between glomeruli. **(F-H)** Tissue from NZC organoid graft. **(F)** Example of NZC organoid at the time of engraftment. **(G)** WT1 staining for podocytes reveals abundant glomeruli. **(H)** CD31 staining for endothelial cells shows vascularization of the graft tissue. **(I)** PDGFRβ staining for stromal cells shows that these are abundant in the graft tissue.

## DISCUSSION

This study shows that heterochronic mixing of cells derived by directed differentiation strongly potentiates the differentiation of kidney tissue in organoids derived from pluripotent stem cells. Nephron progenitor cell contribution to the nascent nephron of the developing kidney is asynchronous (Lindstrom et al., 2018a), and mixing kidney progenitor cells at distinct stages of differentiation is intended to emulate this process. The finding that the two different cell batches in a heterochronic mix contribute differentially to proximal and distal compartments of the nephron supports the idea that heterochronic mixing does indeed promote nephrogenesis by establishing a system for time-dependent differentiation. Further support is based on the observation that the majority of cells derived from directed differentiation undergo conversion to the LHX1-expressing epithelial cell precursor state during the first 2 days after aggregation, and we infer from this that the NPC population is severely depleted. Supplementing with a newly differentiated batch of NPCs at this point replenishes this population, contributing to the proximal nephron regions.

Engrafted organoids differentiated using the heterochronic mixing strategy are efficiently vascularized and form characteristic nephron components in vivo. These are exclusively derived from the engrafted cells with the exception of the endothelium, which appears exclusively derived from the host. This feature of the engraftment is puzzling, since a well-developed network of endothelial cells is found in the organoid prior to engraftment. Since the complex glomerular vasculature can be derived from the host, this finding does not present a limitation, but rather raises the question why endothelial cells differentiated using our modification of the (Morizane et al., 2015) protocol do not contribute to blood vessels in the graft. Endothelial cells are known to be extremely fastidious in their growth requirements and protocols for differentiation of cells with potential to form functional vasculature in vivo differ significantly from the conditions that we used. Thus, it seems likely that our culture conditions do not provide key signals for differentiation of functional endothelial cells.

In vivo analysis of engrafted kidney tissue is essential for longitudinal studies, for example drug treatments. However, simple technology to detect and track subcapsular grafts without sacrificing the host animals has not been available. The use of microCT with iodinated contrast provides a rapid and relatively simple method for evaluating whether grafts have become vascularized and if they are functionally concentrating contrast agent. In its current form, weekly scanning provides information on whether grafts have taken, their size and if they are displaying even enrichment of the contrast agent. In large studies comparing organoids generated from different patient iPSCs or iPSC with different genetic mutations, it will be important to interpret end-point analyses in the form of histology and functional measurements such as FITC-dextran incorporation in light of the dynamic changes in the graft over the engraftment period.

Stromal cells expressing appropriate molecular markers are found between epithelial structures in organoids prior to engraftment. However, this population expands excessively following engraftment. One possible interpretation would be that cells derived from directed differentiation are not stable in their differentiation state, leading to de-differentiation of epithelial cells to stroma. Genetic studies have for example shown that loss of expression of the transcription factor PAX2 causes NPCs to assume a stromal-like identity in the developing kidney (Naiman et al., 2017). Our finding that wild type embryonic kidneys engrafted under the kidney capsule show similar stromal accumulation suggests that this is not a feature of the directed differentiation process, but instead is associated with the micro-environment. One obvious difference between a subcapsular graft and a developing kidney is of course the collecting duct system. To date we have not observed any tubular connections between the graft and the host and based on this we suggest that degenerative changes in the graft are most likely due to the lack of urine outflow. Our future developments of engraftment technology will focus on establishing connections for urine outflow so that we can reach the goal of functional testing in animal disease models.

## MATERIALS & METHODS

### Regulatory compliance

Animal experiments were reviewed and approved by the Institutional Animal Care and Use Committees of Maine Medical Center and the University of Texas Southwestern Medical Center.

### Cell culture

Human iPSC line WTC11 (a kind gift from Bruce Conklin, Gladstone Institute of Cardiovascular Disease) was maintained in mTeSR1 (Stemcell Technologies) on Matrigel (Corning) coated six well plates (Corning) in a 37°C incubator with 5% CO_2_. At 70-80% confluence, cells were dissociated with Accutase (Stemcell Technologies) and plated on Matrigel-coated six well plates with mTeSR™1 containing 10 μM Rho kinase inhibitor, Y27632 (EMD Millipore). Y27632 was removed after the first 48 hours and thereafter media was replaced with fresh mTeSR1 every day. Human H9 (purchased from Wicell) or H9-FP ubiquitously expressing miRFP703 fused to histone H2B (a kind gift from Dr. Andrew McMahon, University of Southern California) were maintained in StemFit (amsbio) with 100 ng/ml human FGF2 (R&D) on Geltrex (Thermo Fisher Scientific) coated six well plates. At 70-80% confluence, cells were dissociated with Accutase and plated on Geltrex-coated six well plates with StemFit containing 100 ng/ml human FGF2 and 10 μM Y27632. After 48 hours of culture, medium was replaced with fresh StemFit containing 50 ng/ml human FGF2. After the next 48, medium was replaced with fresh StemFit containing 25 ng/ml human FGF2.

### Directed differentiation and generation of kidney organoids

We differentiated hPSCs to kidney progenitors according to (Morizane et al., 2015). In brief, WTC11 were plated at a density of 1.4×10^4^ cells/cm^2^ in a six well plate. On day three, once cells became approximately 50% confluent, medium was replaced with Advanced RPMI 1640 (Thermo Fisher Scientific), 1× GlutaMAX ™ (Thermo Fisher Scientific), 10 μM CHIR99021 (Reprocell), and 5 ng/ml Noggin (R&D Systems). On Day 4, medium was replaced with Advanced RPMI 1640 containing 10 ng/ml Activin A (R&D) and 1X GlutaMAX ™. On Day 7, medium was replaced with Advanced RPMI 1640 containing 10 ng/ml FGF9 (R&D) and 1X GlutaMAX ™ for the next 2 days. H9 or H9-FP, cells were plated at a density of 1.7×10^4^ cells/cm^2^ in a six well plate. On day three, once cells became approximately 50% confluent, media was replaced with differentiation medium containing Advanced RPMI 1640, 1× GlutaMAX ™ and 8 μM CHIR99021 (Reprocell). From day 4 to day 9, the same protocol was followed as for WTC11. On Day 9 of directed differentiation, differentiated cells were designated “kidney progenitor cells”. Kidney progenitor cells were harvested with TrypLE Express (Thermo Fisher Scientific) and resuspended at a density of 2.5×10^5^ cells/μl in organoid initiation medium containing APEL2 (Stemcell Technologies), 1.5% PFHM-II (Thermo Fisher Scientific), 100 ng/ml FGF9, 100 ng/ml BMP7 (R&D Systems) and 1 μg/ml Heparin (Sigma-Aldrich). 1 ml/well of this medium was added to wells in a 24 well plate and Isopore Membranes (EMD Millipore) were suspended at the surface of the medium to create an air-liquid interface. Resuspended cells were spotted on top of the filter (2 μl/aggregate). Medium was changed every 48 hours or when it turned yellow. All growth factors were removed from the medium 4 days after aggregation and organoids were cultured for a further 5 days with APEL2 containing 1.5% PFHM-II.

### Organoid generation through heterochronic recombination

For heterochronic recombination of hPSCs, directed differentiation was performed on two batches of cells staggered two days apart. On day 9, the first batch of kidney progenitor cells was aggregated at the air-liquid interface as described above. 2 days later, aggregated cells were gently broken into small cell clusters with a 200μl micropipette and mixed with newly differentiated kidney progenitor cells. 1 dissociated aggregate was mixed with 5×10^5^ kidney progenitor cells in 4μl of organoid initiation medium and re-aggregated in 2 aggregates at the air-liquid interface. Medium was changed every 48 hours or when it turned yellow. All growth factors were removed from the medium 4 days after aggregation and organoids were cultured for a further 5 days with APEL2 containing 1.5% PFHM ‐II.

### NZCs Organoid generation and engraftment

NZCs were derived from E17.5 mouse kidneys according to (Blank et al., 2009) and suspended at a cell density 2.5×10^5^ cells per μl medium containing APEL2, 1.5% PFHM-II, 200 ng/ml of FGF9 and 1 μg/ml of Heparin. In a 24 well plate, 1 ml/well of medium was added per well and Isopore Membranes were suspended at the surface of the medium to create an air-liquid interface. NZCs were aggregated on top of the filter by spotting 2 μl per aggregate. Medium was changed every 48 hours or when it became yellow. 5 days after aggregation, all growth factors were removed and organoids were cultured with APEL2 containing 1.5% PFHM-II throughout for a further 4 days and then engrafted under the kidney capsule of adult mice. Mice were sacrificed 3 weeks after engraftment.

### Embryonic mouse kidneys for engraftment

Pregnant dams were sacrificed at E15.5 and kidneys were dissected out of the embryos. Kidneys were then engrafted under the kidney capsules of adult NOD SCID mice (Charles River Laboratories). Graft recipients were sacrificed one month or two months after the engraftment and the kidneys were sectioned for H&E staining and immunostaining.

### Flow cytometry

Kidney progenitor cells were characterized by flow cytometry at the end of day 9 of the directed differentiation protocol and 2 days after aggregation at the air-liquid interface. Cells were incubated with TrypLE Express for 5 min at 37 °C to generate single cell suspensions and strained through a 40 μm mesh (BD Biosciences). They were then fixed and permeabilized using True-Nuclear Transcription Factor Buffer Set (Biolegend) according to the manufacturer’s protocol. Blocking buffer consisting of 5% donkey serum in PBS was added for 15 min, then cells were incubated with primary antibodies Six2, Lhx1 or WT1 for 30 minutes at 4 °C and incubated with corresponding secondary antibodies conjugated with Alexa Fluor 488 or Alexa Fluor 647. Isotype-matched anti-rabbit IgG and anti-mouse IgG served as controls. Flow cytometry on stained cells was performed using MACSQuant (Miltenyi Biotec).

### Engraftment

Organoids generated through heterochronic recombination were engrafted in NSG mice (n=6 per experimental group). In brief, mice were anesthetized with isoflurane and an incision was made in the flank to exteriorize the kidney. A small incision was made in kidney capsule and organoids (n=20) were placed under kidney capsule using fine blunt forceps.

### Micro computed tomography

Optiprep (Sigma) at a dose of 1.8 g/kg was injected retro-orbitally. Ten minutes after Optiprep injection, mice were anesthetized by intraperitoneal injection of a 16 mg/kg xylazine and 105 mg/kg cocktail. 25 minutes after optiprep injection, the torso of each mouse was scanned for 8-10 min at an isotropic voxel size of 76 microns (70kV, 114μA, 400ms integration time) with a vivaCT 40 scanner (Scanco Medical Inc.). Two-dimensional gray-scale image slices were reconstructed into a three-dimensional tomography. Scans were reconstructed between the proximal end of L1 and the distal end of L5. The region of interest (ROI) for each animal was defined based on skeletal landmarks from the gray-scale images. Snapshots were taken at the best orientation to show graft perfusion.

### Mice perfusion

To study the connection between graft vasculature and the systemic circulation, 100μl of 1 μg/μl FITC-labeled Griffonia Simplificata Isolectin B4 (Sigma) was injected into the retro orbital sinus. To evaluate glucose uptake of proximal tubule cells, 500μl of 2 mg/ml Alexa Fluor 488 conjugated Dextran, 10,000 MW (Thermo Fisher Scientific) was injected into the retro-orbital sinus. 30 minutes after injection, mice were anesthetized with xylazine+ketamine and the right atrium of the heart was removed. Mice were then perfused via the left ventricle with 15ml of 4% paraformaldehyde dissolved in PBS followed by 60 ml PBS. Kidneys were collected for analysis after perfusion.

### Immunostaining

Whole mount staining was performed on organoids and 100μm vibratome sections of kidneys carrying grafts. Organoids were fixed in 4% paraformaldehyde/PBS for 15 minutes at room temperature and then permeabilized with 1% Triton X100/PBS for 10 min at 4°C. Blocking buffer consisting of 5% donkey or goat serum was added for 1 hour and then primary antibodies diluted in blocking buffer were added (Table 1) and organoids were incubated at 4°C overnight. They were then washed with PBS for 6 hours at 4°C and incubated with secondary antibodies (Table 1) and 0.01 μg/ml DAPI at 4°C overnight. Organoids were then washed with PBS for 6 hours and mounted in VECTASHIELD antifade mounting medium (Vector Laboratories). Fluorescent images were taken with a confocal microscope (Leica SP8). For quantification, integrated density of fluorescent images was measured using Image J. For each antibody stain, three separate fields from three different specimens were measured, and the mean was plotted with standard error bars. Kidneys with engrafted organoids were pared down so that only a transverse slab of host kidney underlying the graft remained and this tissue was fixed with 4% paraformaldehyde/PBS for 1 hour per mm thickness of the slab at 4°C before vibratome sectioning at 100μm. Immunostaining was performed using the same method as for organoids. For analysis of the embryonic kidney engraftment, kidneys were fixed overnight in 4% paraformaldehyde/PBS at 4 °C, paraffin embedded and sectioned. Sections were subjected to heat-induced epitope retrieval with Tris-EDTA, blocked in PBS with 0.1% Triton X-100 and 5% FBS and incubated overnight with primary antibodies. Secondary antibodies were added for 1h at room temperature. Slides then were mounted with Vectashield and photographed by scanning laser confocal microscopy (Zeiss LSM-700). The primary antibodies used are shown in Table 1.

**Table 1.** List of antibodies/lectins.

| Name | Catalog/clone number | Source | Dilution |
| --- | --- | --- | --- |
| Cdh1 | ab11512 | Abcam | 1:100 |
| Podxl | AF1658 | R&D systems | 1:100 |
| LTL | B-1325 | Vector | 1:200 |
| Brn1 | sc-6028-R | Santa cruz | 1:100 |
| GATA3 | 5852S | Cell signaling Tech. | 1:100 |
| DBA | B-1035 | Vector | 1:200 |
| WT1 | ab89901 | Abcam | 1:50 |
| CD31 | 3528 | Cell signaling Tech. | 1:50 |
| Pdgfb | ab32570 | Abcam | 1:100 |
| Meis1 | 39795 | Active Motif | 1:100 |
| Hnf4a | C11F12 | Cell signaling | 1:100 |
| Lhx1 | 4F2-c | Developmental Studies Hybridoma Bank | 1:200 |
| Emcn | ab45771 | Abcam | 1:100 |
| SMA | C6198 | Sigma | 1:200 |
| THP | J65429 | Alfa Aesar | 1:200 |
| KRT8 | TROMA-1-c | Developmental Studies Hybridoma Bank | 1:200 |
| Renin | ab212197 | Abcam | 1:100 |
| HuNu | MAB1281B | Millipore | 1:50 |
| Collagen I | 600-401-103-0.1 | Rockland | 1:200 |
| Collagen IV | 600-401-106-0.1 | Rockland | 1:200 |
| Laminin | L9393 | Sigma | 1:200 |
| Six2 | 11562-1-AP | Proteintech | 1:100 |
| Donkey anti-Rabbit IgG, Alexa Fluor 488 | A-21206 | Thermo Fisher Scientific | 1:200 |
| Donkey anti-mouse IgG, Alexa Fluor 488 | A32766 | Thermo Fisher Scientific | 1:200 |
| Donkey anti-rat IgG, Alexa Fluor 488 | A-21208 | Thermo Fisher Scientific | 1:200 |
| Streptavidin, Alexa Fluor 488 | S11223 | Thermo Fisher Scientific | 1:200 |
| Donkey anti-Rabbit IgG, Alexa Fluor 568 | A10042 | Thermo Fisher Scientific | 1:200 |
| Donkey anti-Goat IgG, Alexa Fluor 568 | A-11057 | Thermo Fisher Scientific | 1:200 |
| Goat anti-Mouse IgG1, Alexa Fluor 568 | A-21124 | Thermo Fisher Scientific | 1:200 |
| Streptavidin, Alexa Fluor 568 | S11226 | Thermo Fisher Scientific | 1:200 |
| Donkey anti-Rabbit IgG, Alexa Fluor 647 | A-31573 | Thermo Fisher Scientific | 1:200 |
| Streptavidin, Alexa Fluor® 647 | S21374 | Thermo Fisher Scientific | 1:200 |

## ACKNOWLEDGMENTS

We would like to express our gratitude to Dr. Andrew McMahon and Tracy Tran at University of Southern California for sharing the H9-FP cells. The project described was supported by the National Institutes of Health grant number R24 DK106743 to L.O. and T. C. Core facilities support was provided by Maine Medical Center Research Institute core facilities for Molecular Phenotyping, Progenitor Cell Analysis, Microscopy, and Small Animal Imaging. A special thank you to Terry Henderson at the Maine Medical Center Research Institute Small Animal Imaging core for help with Micro-CT measurements. The content is solely the responsibility of the authors and does not necessarily represent the official views of the National Institutes of Health.

## SUPPLEMENTARY FIGURES

**Figure S1:**
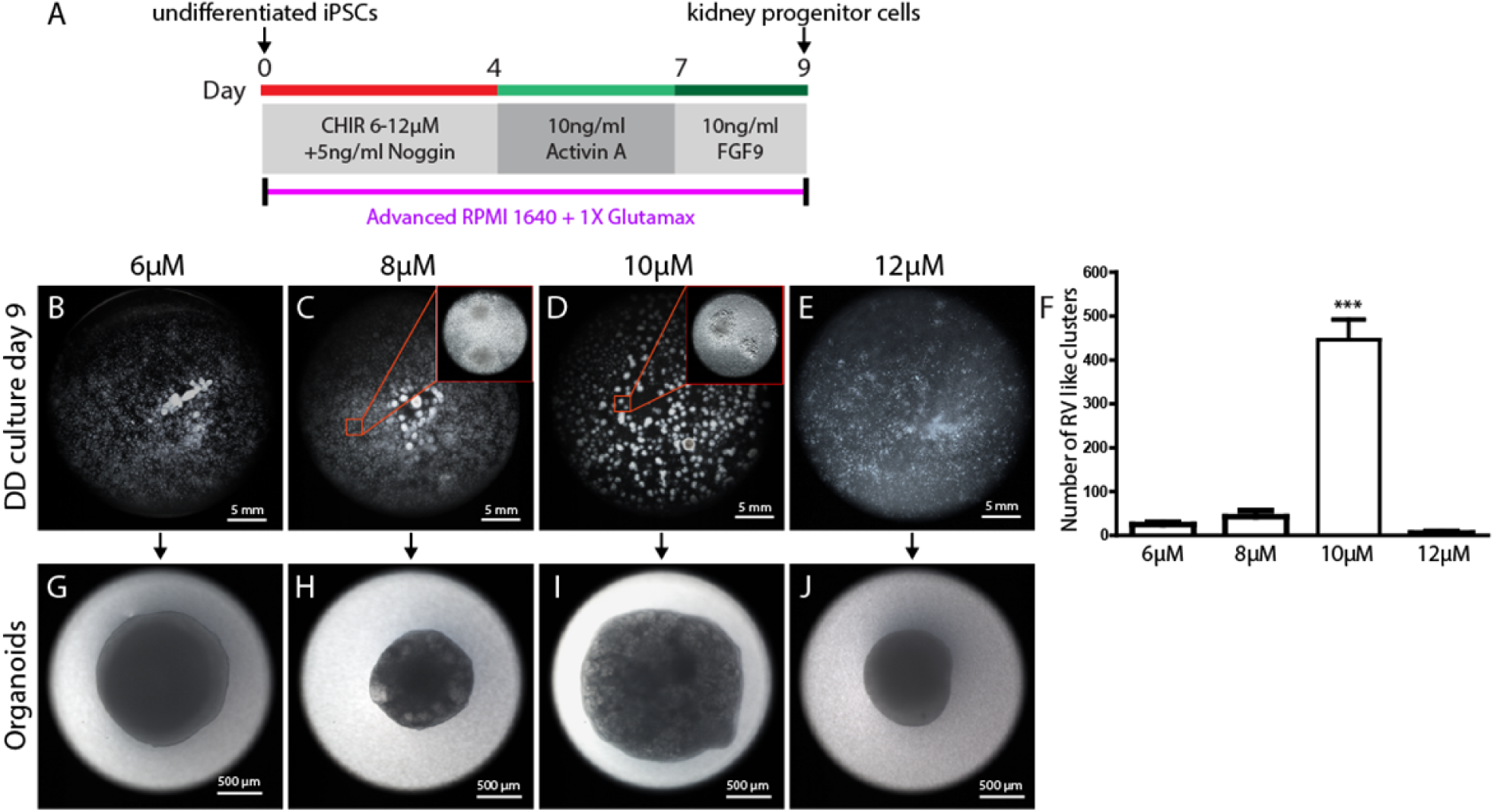
CHIR dose adjustment for directed differentiation of WTC11 human iPSCs. **(A)** The directed differentiation procedure showing the period between initiation of directed differentiation (day 0) and 4 days during which time CHIR was titrated. Stereo microscope images of cell culture plates on day 9 showing clusters of epithelializing cells following treatment with **(B)** 6 μM, **(C)** 8 μM. Inset shows magnified view of sparse cell clusters. **(D)** 10μM. Inset shows magnified view of dense cell clusters with epithelial morphology, and **(E)** 12 μM CHIR. **(F)** Clusters with epithelial morphology were counted. Data obtained from three independent experiments and expressed as mean±SEM, ***p<0.001. **(F-I)** Stereo microscope images showing kidney organoids generated by aggregation and culture at the air-liquid interface as shown in Figure 1 for each directed differentiation.

**Figure S2:**
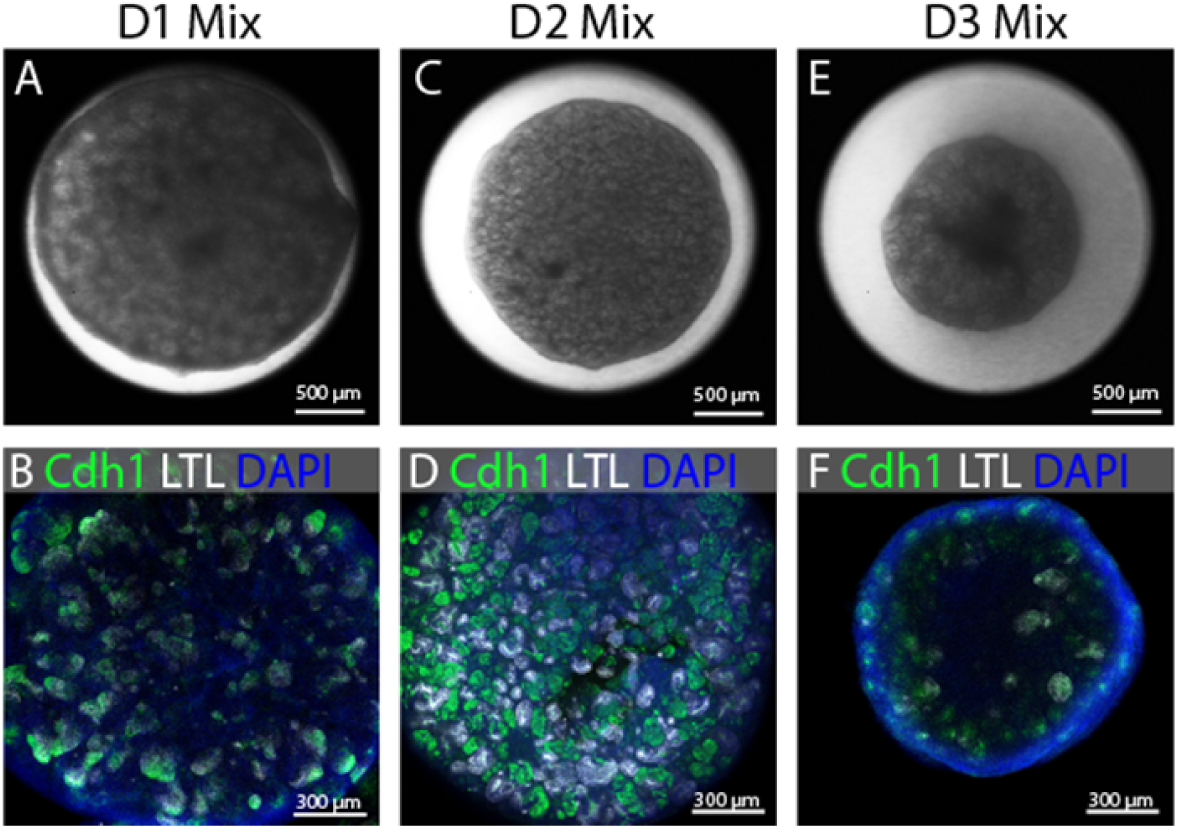
Determination of time point for heterochronic recombination. Kidney progenitors derived from directed differentiation on day 9 were aggregated on air liquid interface and were recombined with a fresh batch of kidney progenitors after 1 day (D1), 2 days (D2) and 3 days (D3). **(A, C and E)** Stereo microscope images showing organoid size and complexity. **(B, D and F)** Immunostaining with Cdh1+ to mark distal epithelia, and LTL to mark proximal. DAPI stains nuclei.

**Figure S3:**
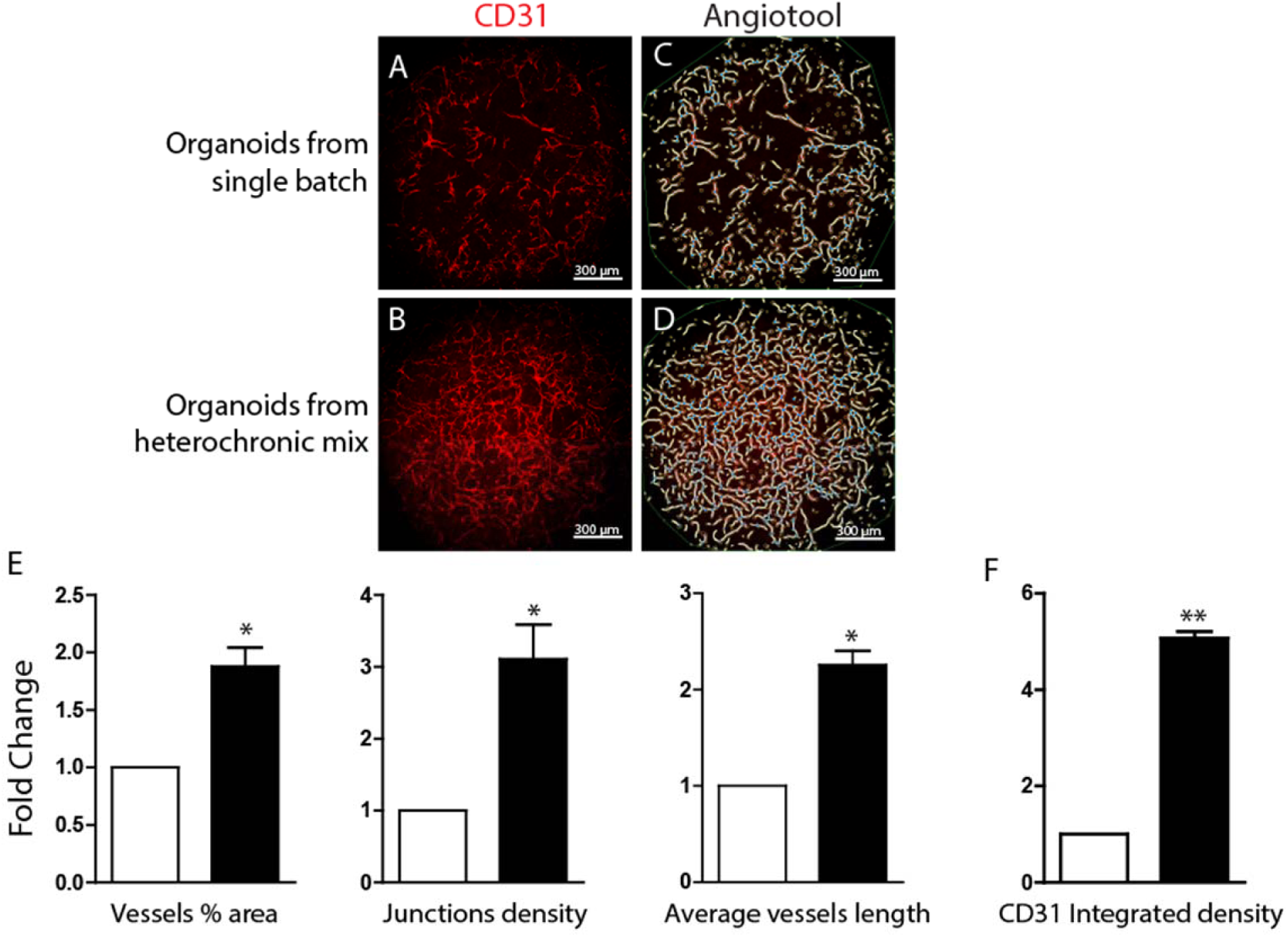
Comparison of endothelial networks in organoids derived from single batches of directed differentiation cells or heterochronic mixes. Immunofluorescence image showing CD31+ endothelial network in organoid generated from **(A)** single batch and **(B)** heterochronic recombination. **(C-D)** Maximal projections of confocal Z-stacks quantified using Angio tool software. White lines depict endothelial networks, blue dots show junctions, yellow lines showing edge of vascular networks and thin green lines showing total area over which organoid was quantified. **(E)** Fold change in vessel % area, junction density and average vessel length was calculated from single mix (open bars) and heterochronic mix organoids (solid bars). **(F)** Fold change in CD31 integrated density was quantified by Image J in single mix (open bars) and heterochronic mix organoids (solid bars). Values calculated from three independent experiments, statistical significance was determined by unpaired t-test and expressed as mean ± SEM; *p<0.05, **p<0.01.

**Figure S4:**
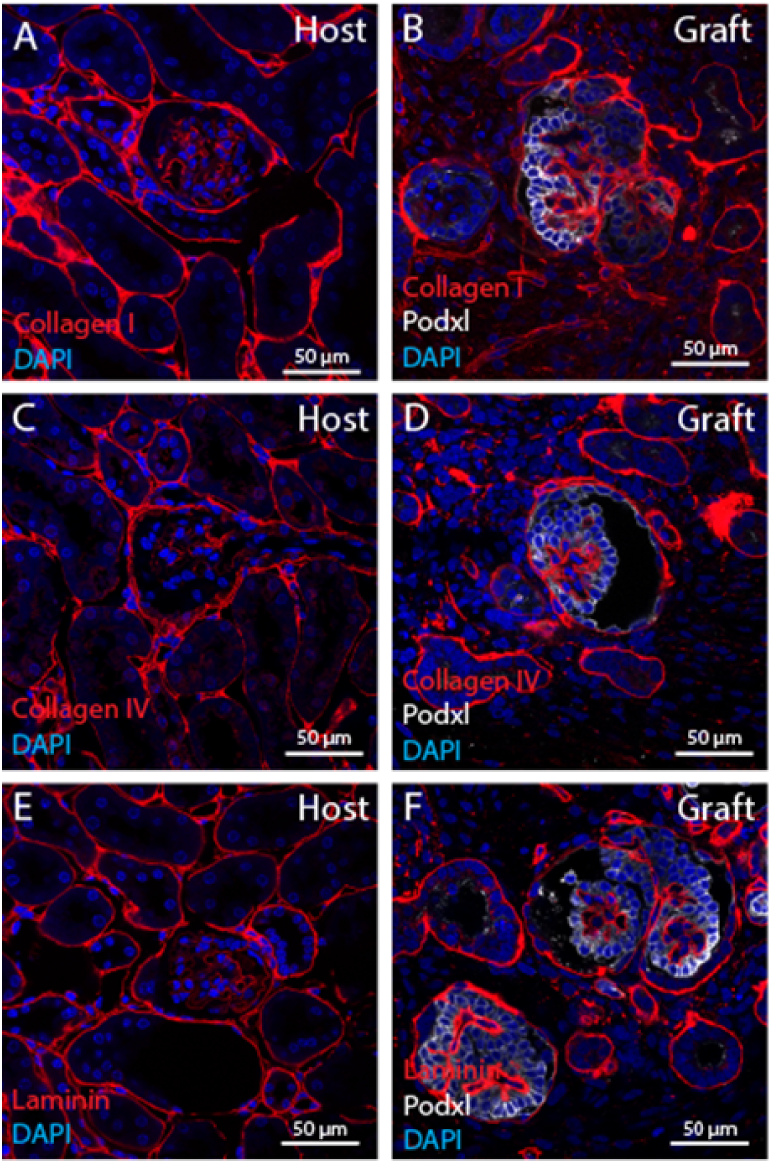
Glomerular basement membrane in the engrafted organoids. (A-C) Engraftment of organoids resulted in more mature nephron structures and presence of Collagen I^+^, Collagen IV^+^ and Laminin^+^ glomerular basement membrane surrounded by Podxl^+^ podocytes in presumptive glomerulus.

